# The ribbon-helix-helix domain proteins CdrS and CdrL regulate cell division in archaea

**DOI:** 10.1101/2020.06.16.153114

**Authors:** Cynthia L. Darnell, Jenny Zheng, Sean Wilson, Ryan M. Bertoli, Alexandre W. Bisson-Filho, Ethan C. Garner, Amy K. Schmid

## Abstract

Precise control of the cell cycle is central to the physiology of all cells. In prior work we demonstrated that archaeal cells maintain a constant size; however, the regulatory mechanisms underlying the cell cycle remain unexplored in this domain of life. Here we use genetics, functional genomics, and quantitative imaging to identify and characterize the novel CdrSL gene regulatory network in a model species of archaea. We demonstrate the central role of these ribbon-helix-helix family transcription factors in the regulation of cell division through specific transcriptional control of the gene encoding FtsZ2, a putative tubulin homolog. Using time lapse fluorescence microscopy in live cells cultivated in microfluidics devices, we further demonstrate that FtsZ2 is required for cell division but not elongation. The *cdrS-ftsZ2* locus is highly conserved throughout the archaeal domain, and the central function of CdrS in regulating cell division is conserved across hypersaline adapted archaea. We propose that the CdrSL-FtsZ2 transcriptional network coordinates cell division timing with cell growth in archaea.

**Importance:** Healthy cell growth and division are critical for individual organism survival and species long-term viability. However, it remains unknown how cells of the domain Archaea maintain a healthy cell cycle. Understanding archaeal cell cycle is of paramount evolutionary importance given that an archaeal cell was the host of the endosymbiotic event that gave rise to eukaryotes. Here we identify and characterize novel molecular players needed for regulating cell division in archaea. These molecules dictate the timing of cell septation, but are dispensable for growth between divisions. Timing is accomplished through transcriptional control of the cell division ring. Our results shed light on mechanisms underlying the archaeal cell cycle, which has thus far remained elusive.

## Introduction

The cell cycle proceeds through an ordered progression of molecular events including cell volume increase, DNA replication, segregation, and cytokinesis. The fine-tuned control between these processes has been studied intensely for decades, yielding deep insight into cell cycle mechanisms. To date, such work has focused on bacterial and eukaryotic model organisms. In contrast, the archaeal cell cycle remains virtually unexplored despite its importance as the evolutionary progenitor of eukaryotes (Zaremba-Niedzwiedzka et al., 2017). The few studies that have been conducted on the cell cycle in archaeal model organisms point to a hybrid of eukaryotic and bacterial features with differential assortment of these features throughout the archaeal lineages. For example, in *Crenarchaeota*, the cell cycle phases, molecular machinery of DNA replication, and cell division are largely conserved with eukaryotes (Kelman and Kelman, 2014; Lindas and Bernander, 2013; Lundgren and Bernander, 2007; Poplawski and Bernander, 1997). In contrast, the cell cycle in the lineage Euryarchaeota retains features of all three domains, including a bacterial FtsZ system of cell division (Makarova and Koonin, 2010), a eukaryotic system for DNA replication (Kelman and Kelman, 2014), and archaeal-specific firing of replication origins (Hawkins et al., 2013).

The tubulin homolog FtsZ has been studied in detail in many bacteria for its central role in cell division. In most bacteria, FtsZ monomers assemble into short filaments that form the cytokinetic ring at mid-cell, which constricts to divide the mother cell into two daughters of equal size (Adams and Errington, 2009; Bisson-Filho et al., 2017; Haeusser and Margolin, 2016; Xiao and Goley, 2016; Yang et al., 2017). FtsZ in archaea appears to function similarly, as previous fluorescence imaging experiments in fixed (Aylett and Duggin, 2017; Grant et al., 2018; Herrmann and Soppa, 2002; Margolin et al., 1996) and live (Abdul-Halim et al., 2020; Duggin et al., 2015; Walsh et al., 2019) hypersaline-adapted archaeal cells (species *Haloferax volcanii*) demonstrated Z-like rings forming at mid-cell. However, all known halophilic archaeal genomes encode multiple tubulin-like proteins (Aylett and Duggin, 2017; Becker et al., 2014), so the function and mechanism of these proteins in cell division remain unclear.

Halobacteria, a clade of hypersaline-adapted Euryarchaeota, provide excellent model systems for understanding cell cycle mechanisms and how they are regulated. In particular, for the model species *Halobacterium salinarum, Haloferax volcanii*, and *Haloferax mediterranei*, large and facile toolkits enable genetic manipulation (knockouts, overexpression, etc.)(Allers et al., 2010; Bitan-Banin et al., 2003; Liu et al., 2011; Peck et al., 2000). For *Hbt. salinarum* strain NRC-1, large systems biology datasets, including transcriptomic profiles under a wide array of growth and stress conditions, enable rapid hypothesis generation regarding gene functions (Brooks et al., 2014; Dulmage et al., 2018). In previous work, we developed live-cell, time-lapse microscopy methods for hypersaline adapted archaea to overcome the challenges of rapid salt crystallization on microscopy slides (Eun et al., 2018). Salt-impregnated agarose microchambers were fabricated using soft lithography, which support up to six generations of growth for *Hbt. salinarum.* Using these tools, we demonstrated that single, rod-shaped *Hbt. salinarum* cells grow (elongate) exponentially, adding a constant volume between divisions (the “adder” model of cell size control (Sauls et al., 2016)). However, the size distribution and division site placement at mid-cell demonstrated greater variance than bacterial cells that maintain their size in a similar fashion (Eun et al., 2018). Here we adapt microfluidics for *Hbt. salinarum* and leverage the existing genetics and systems biology toolkits to interrogate the regulation of the archaeal cell cycle.

Cell cycle progression in eukaryotes is known to be exquisitely regulated, and DNA replication and cell division are coordinated in bacteria (Haeusser and Levin, 2008). However, despite recent progress regarding cell growth and size control in archaea, the underlying molecular mechanisms regulating these processes remain unknown. Gene expression profiling experiments suggest that archaea possess the capability for oscillating gene expression patterns, a hallmark of genes with cell cycle-related functions in eukaryotes (Orlando et al., 2008). For example, our prior work with transcriptomics in *Hbt. salinarum* provides evidence for temporally coordinated induction of hundreds of genes during the resumption of growth following stasis (Schmid et al., 2007; Skotheim et al., 2008). Oscillating gene expression was observed in *Hbt. salinarum* cultures entrained to day-night cycles (Whitehead et al., 2009). Cyclic gene expression patterns have also been observed in synchronized cultures of the crenarchaeon *Sulfolobus solfataricus* (Lundgren and Bernander, 2007).

Gene regulatory networks (GRNs), comprised of interacting transcription factors (TFs) and their target genes, are central to the process of dynamic, physiological response to a variable environment. Archaeal transcription proteins resemble those of both bacteria and eukaryotes at the level of amino acid sequence. Basal transcriptional machinery required for transcription initiation in archaea, like that of eukaryotes, consists of transcription factor II B (TFB), a TATA binding protein (TBP), and an RNA-Pol II-like polymerase (reviewed in (Martinez-Pastor et al., 2017)). The proteins that modulate transcription (*e.g.,* activator and repressor TFs) typically resemble those of bacteria, with the majority of these proteins possessing HTH or wHTH DNA binding domains (Perez-Rueda and Janga, 2010). Our recent studies on gene regulatory networks (GRNs) in *Hbt. salinarum* systematically investigated the function of transcription factors using high throughput phenotyping of TF knockouts (Darnell et al., 2017; Tonner et al., 2017). This study implicated the putative TF DNA binding protein VNG0194H (VNG_RS00795) as a candidate regulator of multiple stress responses: deletion of *VNG0194H* lead to growth defect under multiple stress conditions, including oxidative stress, low salinity, and heat shock (Darnell et al., 2017). Intriguingly, *VNG0194H* is encoded upstream of *ftsZ2* (Ng et al., 2000), suggesting additional roles for VNG0194H in cell growth and/or division. An additional putative DNA binding transcriptional regulator VNG0195H is encoded upstream.

To address knowledge gaps regarding archaeal cell division mechanisms, here we investigate the cell growth and division functions of FtsZ2, VNG0194H (CdrS, cell division regulator short) and VNG0195H (CdrL, cell division regulator long). We combine a battery of assays, including genetic knockouts, quantitative time lapse microscopy of single cells, custom microfluidics technology, gene expression profiling, and TF-DNA binding ChIP-seq experiments. The resultant data demonstrate that CdrS and FtsZ2 are required for normal cytokinesis but not cell elongation. This regulation is accomplished via: (i) CdrS activation of *ftsZ2* and other cell cycle-related genes; (ii) CdrL direct regulation of the *cdrS-ftsZ2* operon. The CdrSL GRN system is highly specific to regulation of *ftsZ2* at the level of transcription.

## Results

### *cdrS* encodes a conserved, putative transcription factor co-transcribed with the tubulin-encoding *ftsZ2* gene

Our previous genetics experiments indicated an important role for putative DNA binding protein VNG0194H in stress response and growth physiology of *Hbt. salinarum* (Darnell et al., 2017; Tonner et al., 2017). We first used bioinformatics to generate hypotheses regarding the physiological function of VNG0194H and its encoding gene locus (Figure 1A)*. VNG0194H* is predicted to encode a small 55 amino acid, single-domain protein that exhibits >99% structural homology to other ribbon-helix-helix domain transcriptional regulators of the RHH_1 family (PF01402, E-value of primary sequence homology 5.3 × 10^−5^; 99.6% confidence in structural homology to *E. coli* NikR), suggesting that it may function as a DNA binding transcriptional regulator or in protein-protein interactions (Gomis-Ruth et al., 1998). Encoded immediately upstream, the VNG0195H protein is predicted to contain an N-terminal RHH_1 domain (95.9% confidence in structural homology to *E. coli* NikR) and a C-terminal double zinc ribbon domain (DZR, PF12773, E-value 3.1×10^−7^). The presence of VNG0195H in this locus appears to be unique to halophiles (Supplementary Figure S1). In addition, a tubulin FtsZ homolog is encoded immediately downstream of VNG0194H (*ftsZ2*; *VNG0192G;* Figure 1A). *Hbt. salinarum* FtsZ2 exhibits strong primary sequence identity to known tubulin components of the cell division ring (tubulin / FtsZ GTPase domain PF00091, E-value 7.4×10^−77^; tubulin C-terminal domain PF03593, E-value 6×10^−32^; (Duggin et al., 2015)). Taking these bioinformatic analyses together, we re-named VNG0194H as “CdrS” for “cell division regulator Short” and VNG0195H as “CdrL” for “cell division regulator Long”.

**Figure 1.**
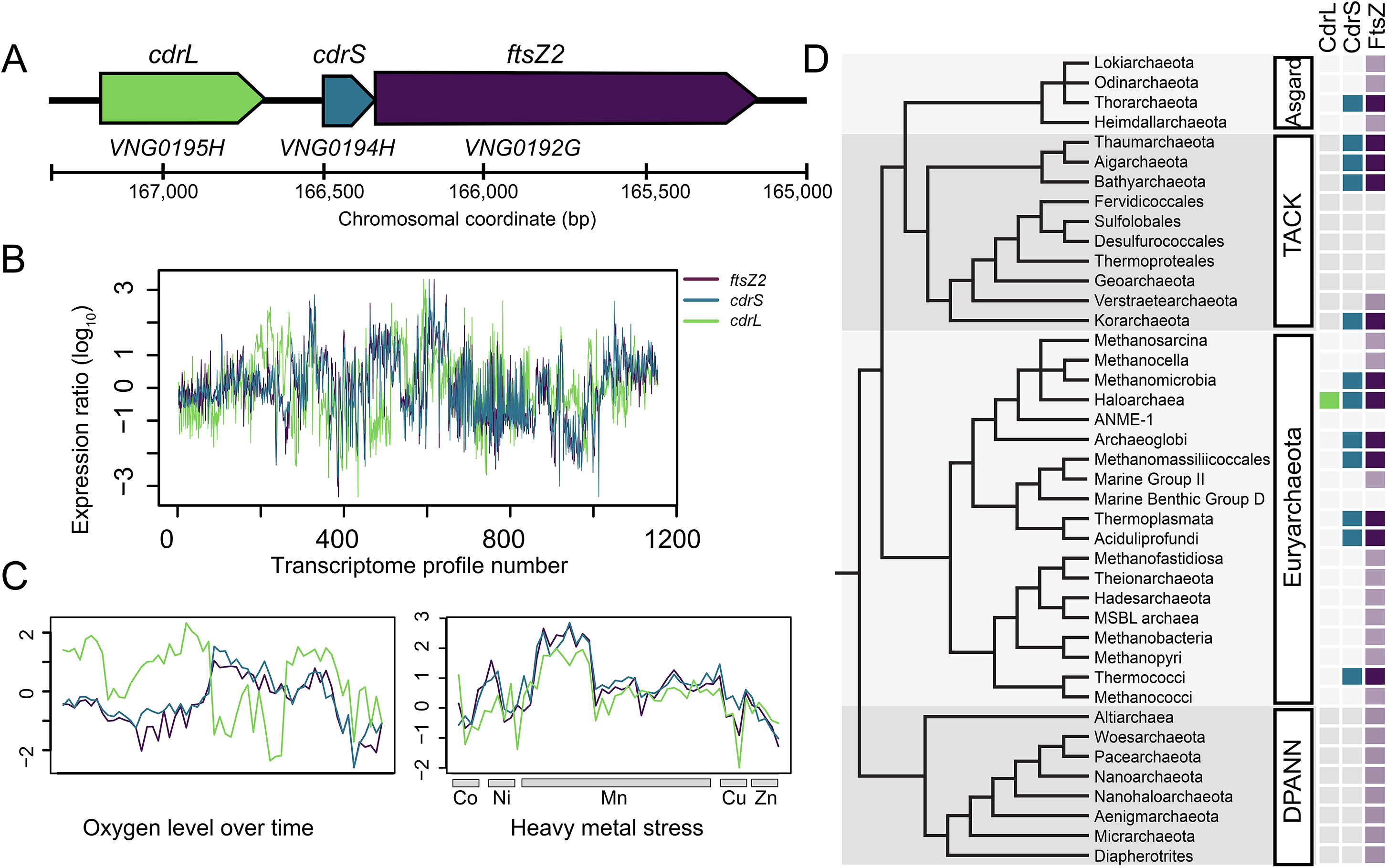
*cdrS* and *ftsZ2* comprise a polycistronic locus conserved throughout archaeal genomes. (A) Locus organization of *cdrL*, *cdrS*, and *ftsZ2* genes (*VNG_RS00800* – *VNG_RS00790; VNG0195H-VNG0192G*). Genes are drawn to scale, with chromosomal coordinates and scale bar shown below the locus diagram. (B) log10 gene expression data from 1,154 normalized microarray experiments over various conditions and genetic backgrounds. X-axis indicates the transcriptome profile number across multiple growth and stress conditions (Brooks et al., 2014; Dulmage et al., 2018). Y-axis represents normalized log_10_ gene expression ratio relative to the wild type control under optimum growth conditions (mid-logarithmic phase, rich CM medium, 37°C, 225 rpm shaking). (C) Gene expression under subsets of conditions in which *cdrL* is not co-expressed (left) or is co-expressed (right) with *cdrS* and *ftsZ2*. Axes are as in panel C. (D) Co-occurrence of *ftsZ2* and *cdrS* in the major archaeal lineages (Hug et al., 2016; Raymann et al., 2015; Spang et al., 2017). Light purple boxes indicate clades with genomes found to encode at least one FtsZ protein but no CdrS (see also Supplementary Table 1). Blue and dark purple boxes indicate clades with genomes that encode genes homologous to *cdrS* and *ftsZ2* in synteny (see also Supplementary Figure S1).

Previous tiling microarray studies indicated co-transcription of *cdrS* and *ftsZ2*, but were inconclusive regarding the inclusion of *cdrL* in this operon (Koide et al., 2009). To further investigate the transcriptional status of this locus, we examined the expression of the three genes from 1,154 microarray and 3 RNA-seq transcriptome profiles for *Hbt. salinarum* grown under a wide variety of environmental and genetic perturbations (Brooks et al., 2014; Dulmage et al., 2018)(Figure 1B). Across all conditions, *cdrS* and *ftsZ2* were strongly and significantly correlated (Spearman’s ρ = 0.923, 95% CI = 0.915, 0.932), whereas *cdrL* was weakly but significantly correlated with *cdrS* (ρ = 0.275, 95% CI = 0.221, 0.327) and *ftsZ2* (ρ =0.299, 95% CI = 0.245, 0.350). As a control, we calculated the correlation of these genes with an unrelated gene located elsewhere in the genome (*trmB VNG1451C*), which exhibited weakly negative correlation with the locus (*cdrS*, ρ = −0.275, 95% CI = −0.327, −0.221; *ftsZ2*, ρ = −0.291, 95 % CI = −0.343, −0.238). The weak but significant correlation of *cdrL* with *ftsZ2* and *cdrS* is driven by strong co-expression of the three genes in response to stress such as metal overload (50 transcriptome profiles, ρ = 0.771, CI = 0.627, 0.864; Figure 1C, right). In contrast, *cdrL* is not co-expressed with *ftsZ2* or *cdrS* under conditions that foster rapid growth (58 profiles; ρ = −0.158, 95% CI = −0.400, 0.104; Figure 1C, left). Previous statistical models that inferred the global gene regulatory network of *Hbt. salinarum* also predicted co-regulation of *cdrS* and *ftsZ2* under all growth conditions, whereas *cdrL* was only co-regulated with the other two genes under a subset of conditions (Brooks et al., 2014). Together these results suggest that *cdrS* and *ftsZ2* are co-transcribed from a polycistronic operon that is co-regulated under all growth conditions. *cdrL,* in contrast, is conditionally co-regulated with the other two genes.

To determine how broadly this locus was conserved outside of *Hbt. salinarum*, we investigated the sequence conservation of each gene and the genomic synteny of the gene pair across archaeal genomes. The co-occurrence of *cdrS* with *ftsZ2* homologs was detectable across representatives of all known archaeal clades except DPANN (Figure 1D; Supplementary Table 1. All supplementary files available at https://doi.org/10.6084/m9.figshare.12195081.v1). FtsZ in the absence of CdrS was also widely distributed. Conservation of the *cdrS-ftsZ2* locus was particularly strong across the Euryarchaeota, with wide conservation across the halophilic archaeal clade (including classes Halobacteria, Natrialbales, and Haloferacales) and neighboring phylogenetic class Methanomicrobia (Supplementary Figure S1). CdrL was widespread across the Halobacteria but absent from all other archaeal clades. Taken together, these results suggest that: (a) *cdrS* exhibits a strong primary and secondary structural homology to transcriptional regulators of the CopG family; (b) the *cdrS-ftsZ2* locus encodes a highly conserved, co-regulated transcriptional unit; (c) CdrL is a putative transcription regulator unique to halophiles and appears to be conditionally co-expressed with *cdrS-ftsZ2*. We therefore focused our subsequent analysis on CdrS.

### The *cdrS-ftsZ2* locus is important for maintaining cell size and biomass in bulk culture

To test the function of the *cdrS-ftsZ2* locus, we constructed independent gene deletion mutants in each coding region and tested for cell division defects. We previously reported a Δ*VNG0194H* strain (Darnell et al., 2017; Tonner et al., 2017); however, that strain included a start site for the *ftsZ2* gene that was mis-annotated in the NCBI database, which we have corrected here (Figure 1A, Supplementary Figure S1). This enabled a more precise, conservative deletion within *cdrS* to avoid polar effects by keeping the putative *ftsZ2* ribosome binding site intact (Figure 1A, Supplementary Tables 5-7). Because halophilic archaea are highly polyploid (Zerulla and Soppa, 2014), stringent quality controls were implemented for both Δ*cdrS* and Δ*ftsZ2* strains, including PCR, Sanger sequencing, and whole genome Illumina re-sequencing (see Methods). These controls confirmed that the coding genes were removed from all genome copies and that no second site mutations had accumulated (Supplementary Table 2).

To investigate the phenotypes of the Δ*cdrS* and Δ*ftsZ2* mutant strains, cells were grown in aerobic batch culture in rich medium. Population growth rates, colony forming units (CFU), and single cell length and area were quantified (Methods). Early exponential phase cultures of the Δ*ura3* parent strain were comprised of cells with a mean length of 5.39 μm (σ = 3.076 μm; Table 1; Supplementary Table 3) and an area of 6.32 μm^2^ (σ = 4.12 μm^2^; Figure 2A, top panel). 87.9% of Δ*ura3* cells fell within one standard deviation of the geometric mean for length (Figure 2A, insets; Table 1). These length and area measurements are consistent with previous observations for *Hbt. salinarum* growth and division (Eun et al., 2018). The rightward skew of the distribution resulted in 10.3% of the cells longer than one standard deviation above the geometric mean; these longer cells are commonly seen during routine culturing and contribute to the noise in the *Hbt. salinarum* cell division model (Eun et al., 2018). Similar cell lengths were measured at time points sampled in mid-log phase and stationary phases of the growth curve (Table 1; Supplementary Figure S2). In contrast, early exponential phase cultures of the Δ*cdrS* strain cells were significantly longer (Welch’s *p* < 2 × 10^−16^; medium effect size 0.760) and larger in area (*p* < 2.67 × 10^−16^; medium effect 0.665) than those of the parent strain. Large variation in size distribution of mutant length and area were also detected (Table 1; Figure 2A, middle panel; Supplementary Table 3). Indeed, 36% of Δ*cdrS* cells were longer than those of the parent strain, with the longest cells > 40 μm. Similarly, the Δ*ftsZ2* strain cell size was significantly larger and more variable than that of the parent strain (length *p* < 2.2 × 10^−16^, large effect 0.832; area *p* < 3.52 × 10^−15^, medium effect 0.571; Figure 2A, bottom panel). These differences in cell sizes between the parent and mutant strains were consistent across growth phases. Together, these microscopy results indicate a role for *ftsZ2* and *cdrS* in maintaining wild type cell size in *Hbt. salinarum*.

**Figure 2.**
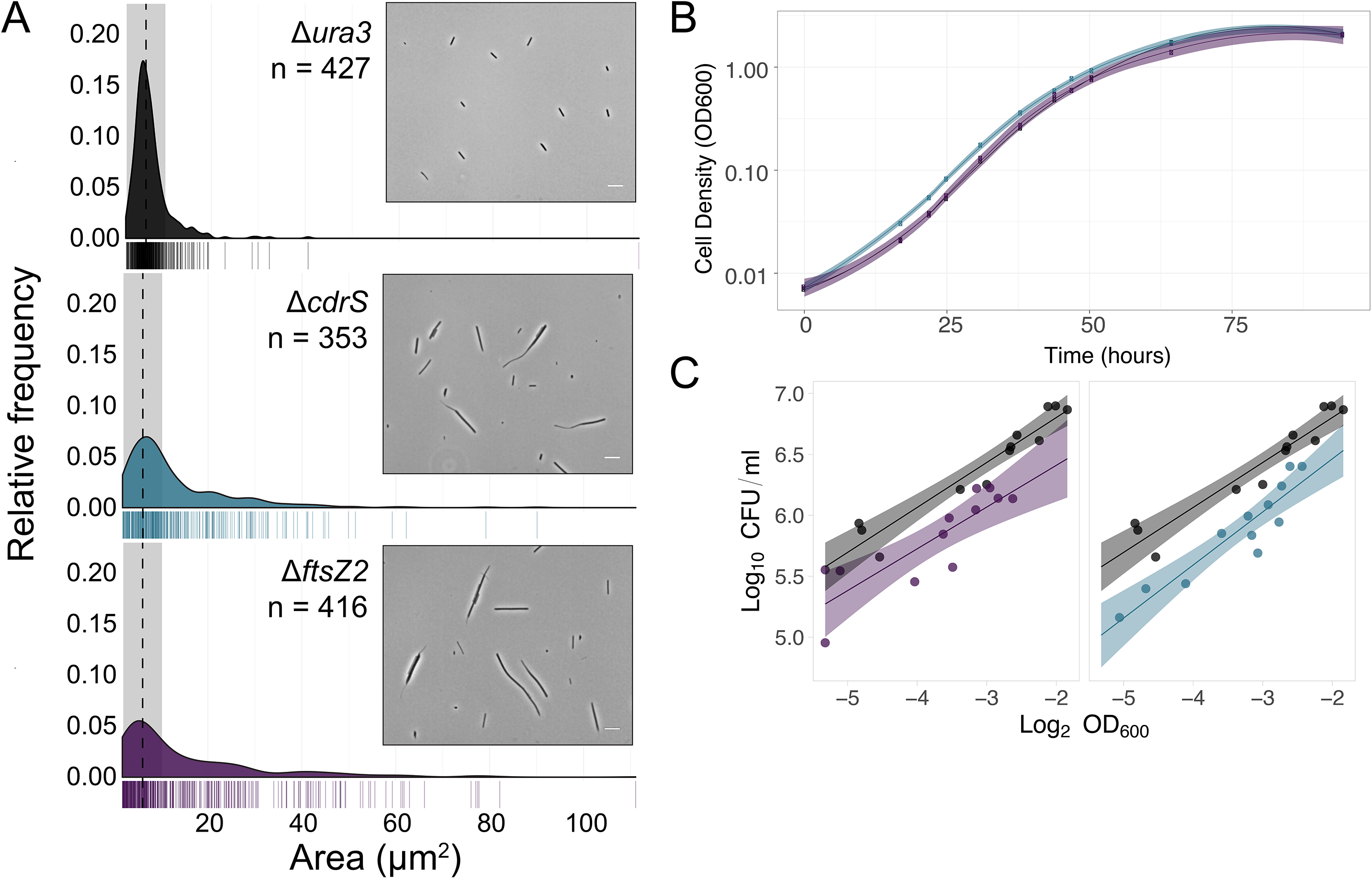
CdrS, and FtsZ2 are required for maintaining cell size and division in batch culture but are dispensable for growth rate. (A) Area of individual cells during early exponential phase across Δ*ura3* (black), Δ*cdrS* (blue), and Δ*ftsZ2* (purple) strains. Median area for Δ*ura3* is indicated in black dashed line. Gray shading indicates 1 standard deviation flanking the Δ*ura3* median in both directions. Insets: Representative phase-contrast micrographs of the Δ*ura3* parent strain and mutants during early exponential growth. White scale bar is 10 μm. (B) Growth curve for all strains in rich media with aerobic conditions measured using optical density at 600 nm. Solid lines represent the mean of three independent biological replicate samples and shaded regions represent 95% confidence intervals. Black, Δ*ura3*; blue, Δ*cdrS*; purple, Δ*ftsZ2*. (C) Correlation of cell concentrations by CFUs per ml and OD_600_. Dots in each panel represent quantification at multiple time points sampled from three replicate exponentially growing batch cultures for each strain. Solid lines indicate the linear regression fit to the data points with shaded regions representing 95% confidence interval. Colors are consistent with panels A and B.

**Table 1.** Cell area geometric means during batch culture (μm^2^ ± σ).

|  | growth phase | early exponential | mid-exponential | early stationary | stationary |
| --- | --- | --- | --- | --- | --- |
| length | $\Delta\text{ura3}$ | $5.39 \pm 3.08$ | $5.72 \pm 3.15$ | $5.27 \pm 2.76$ | $5.09 \pm 2.99$ |
| | $\Delta\text{cdrS}$ | $8.67 \pm 8.70$ | $7.52 \pm 8.02$ | $5.23 \pm 5.48$ | $5.48 \pm 4.15$ |
| | $\Delta\text{ftsZ2}$ | $9.56 \pm 10.04$ | $7.65 \pm 9.46$ | $5.75 \pm 6.52$ | $5.24 \pm 4.65$ |
| area | $\Delta\text{ura3}$ | $6.33 \pm 4.12$ | $6.73 \pm 5.48$ | $5.94 \pm 4.42$ | $6.21 \pm 4.90$ |
| | $\Delta\text{cdrS}$ | $9.50 \pm 11.74$ | $8.97 \pm 16.31$ | $6.16 \pm 9.19$ | $6.40 \pm 7.53$ |
| | $\Delta\text{ftsZ2}$ | $9.76 \pm 16.09$ | $7.18 \pm 14.12$ | $6.14 \pm 10.90$ | $5.64 \pm 8.64$ |

**Table 2.** Geometric means of cell lengths during aphidicolin addition and removal.

| Strain | Time 0 | 6 h post drug addition | 11 h post drug removal |
| --- | --- | --- | --- |
| $\Delta\text{ura3}$ | $4.42 \pm 2.77$ | $7.44 \pm 3.81$ | $4.31 \pm 3.63$ |
| $\Delta\text{cdrS}$ | $8.73 \pm 9.05$ | $9.43 \pm 8.55$ | $6.53 \pm 7.07$ |
| $\Delta\text{cdrS ura3}::P_{\text{cdrS}}\text{-cdrS}$ | $6.45 \pm 7.68$ | $9.52 \pm 7.42$ | $4.77 \pm 5.54$ |

However, it remained unclear whether the drastic increase in cell size together with morphology defects in Δ*ftsZ2* and Δ*cdrS* mutants were due to unregulated growth (cell elongation or biomass accumulation) or a decrease in cell division (fewer septation events). To compare the rate of biomass accumulation between wild type and mutant strains, we measured growth rates by optical density for each of the parent, Δ*ftsZ2*, and Δ*cdrS* strains in batch culture under aerobic conditions in rich media (Methods). The maximum instantaneous growth rate of the Δ*ura3* strain was 0.152 h^−1^ ± 0.004 (Supplementary Table 3), consistent with previous observations (Darnell et al., 2017; Tonner et al., 2017). Neither the Δ*cdrS* or Δ*ftsZ2* mutant strain exhibited a growth rate defect measured by optical density at 600 nm (OD), and reached similar carrying capacities (OD600 = 1.73 – 2.62 for all three strains after 94 hours, Supplementary Table 3, Figure 2B). This suggests that deletion of neither *cdrS* nor *ftsZ2* reduces biomass as measured by optical density.

However, in the spectrophotometer, elongated cells scatter light differently than short cells (Stevenson et al., 2016), which can obfuscate true defects in cell size and/or division. Therefore, in addition to OD readings, we also plated for colony forming units (CFUs). As the largest range of cell lengths occurred during early exponential phase, we plated multiple timepoints in the linear OD range between lag phase and OD = 0.2. We detected a strong and significant positive correlation between log_2_-transformed OD and log_10_-transformed CFUs/ml for the Δ*ura3* parent strain (Pearson’s ρ = 0.9561, *p*-value = 1.188 × 10^−6^; Figure 2C). Similar correlations were detected for each of Δ*cdrS* (ρ = 0.8166, *p*-value = 1.19 × 10^−3^) and Δ*ftsZ2* (ρ = 0.8509, *p*-value = 4.494 × 10^−4^). These strong correlations enabled direct comparison between strains of the CFUs normalized by OD (see Methods). The Δ*ftsZ2* strain yielded 2.2-fold fewer CFUs/ml per log_2_(OD) compared to the Δ*ura3* parent, and Δ*cdrS* had 2.6-fold fewer. These results suggest that fewer viable individual cells are present in cultures of the mutant strains compared to Δ*ura3*, which is likely the result of larger mutant cell size (more biomass per CFU). Together, these colony counts, cell density, and quantitative microscopy results support the hypothesis that *cdrS* and *ftsZ2* gene products are important for cell division but not cell area increase in batch culture.

### Δ*cdrS* cells are phenotypically insensitive to aphidicolin

To further test whether growth and cell division are decoupled in the Δ*cdrS* mutant, we synchronized populations of cells by treating with the cell cycle inhibitor aphidicolin, which specifically targets DNA polymerase-α in eukaryotes (Ikegami et al., 1978). Aphidocolin has been shown to impair DNA replication and cell division but not elongation in wild type *Hbt. salinarum* (Forterre et al., 1984; Herrmann and Soppa, 2002; Schinzel, 1984). Here we quantified cell area prior to aphidicolin addition, following 6 hours of cell cycle block in the presence of drug, and 11 hours after drug removal (Figure 3). After aphidicolin treatment, the Δ*ura3* average cell area increased significantly from 4.42 μm^2^ to 7.44 μm^2^ (geometric mean; Figure 3A; *p* < 2.0 × 10^−16^; large effect size (es) of 1.146), suggesting continued elongation in the absence of division and consistent with previous observations (Herrmann and Soppa, 2002). After aphidicolin removal, Δ*ura3* cells returned to an average area indistinguishable from that of pre-treatment values (4.31 μm^2^; *p* = 0.59; large es 0.965). Cells undergoing septation were also observed, indicating a recovery of cell division (Supplementary Figure S3). In contrast, the distribution of Δ*cdrS* cell area remained largely unchanged by aphidicolin treatment (geometric mean of 8.73 μm^2^ before addition and 9.43 μm^2^ after 6 hours of treatment, respectively; Figure 3B; *p* = 0.15; negligible es 0.094). Removal of aphidicolin from Δ*cdrS* cultures slightly decreased the cell area relative to the pre-treatment area (6.52 μm^2^; 2.3 × 10^−7^; small es 0.433); however, it is unclear whether this decrease is a biological or technical effect, as longer cells may shear during wash steps. Nevertheless, the distribution of cell area in the Δ*cdrS* mutant remained heavily skewed toward elongated cells compared to the Δ*ura3* parent.

**Figure 3.**
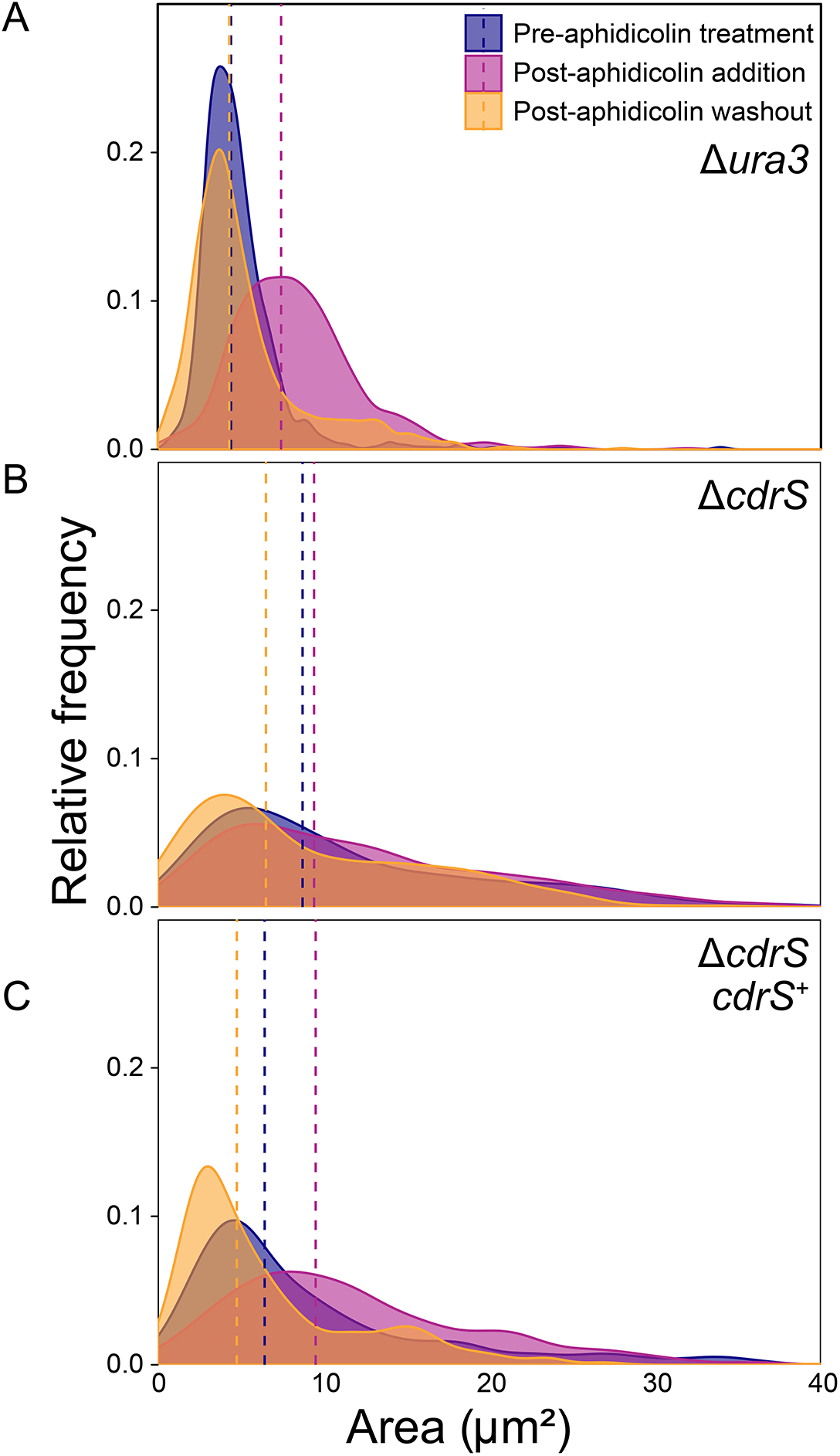
Δ*cdrS* is insensitive to cell cycle inhibitor aphidicolin. Cell area distributions are shown for the Δ*ura3* parent strain (A), Δ*cdrS* strain (B), and the complementation strain (C). As shown in the legend: before drug addition (blue), after 6 hours of exposure to aphidicolin (pink), and 11 hours after removing drug by washing (orange). Dotted lines indicate the geometric mean for each timepoint.

To ensure that the insensitivity to aphidicolin was specific to the deletion of *cdrS*, we generated a complementation strain by integration of *P*_*cdrS*_-*cdrS* into the chromosome at a neutral locus (NC_002607.1:1245981-1247318; Peck et al 2000; Methods). Prior to aphidocolin treatment, the complemented strain cell area intermediate between that of the Δ*ura3* parent and Δ*cdrS* mutant strains (6.45 μm^2^), suggesting partial complementation (Figure 3C). However, after exposure to aphidicolin, the average cell area increased to 9.52 μm^2^ (*p* < 3.9 × 10^−14^, small es 0.448). After aphidicolin wash-out, the cell area mean returned to a smaller 4.78 μm^2^ (2 × 10^−16^, large es 0.887). These shifts in distribution across the time course indicate that the complemented strain was responsive to aphidicolin, suggesting that complementation was achieved under native transcriptional control at the ectopic site. Therefore, while the area of the Δ*ura3* and the *cdrS* complementation strain were affected by aphidicolin treatment, the Δ*cdrS* mutant remained insensitive. Consistent with results from batch culture growth, these data suggest that CdrS is important for cell division but not elongation. In addition, these data support the idea that *cdrS* may act as a checkpoint regulator, acting the pathway that arrests cell division when DNA replication is perturbed.

### Time lapse microscopy reveals cell division defects in Δ*cdrS* and Δ*ftsZ2* mutants

In previous workxs, we designed agarose chambers that allowed real-time microscopic observation of unperturbed growth of *H. salinarum* (Eun et al., 2018). However, these chambers were square in shape (10 × 10 μm), precluding growth of the filamentous Δ*ftsZ2* and Δ*cdrS* strains and necessitating a new device. We adapted the mother machine microfluidic device for real-time growth observation of these mutant *Hbt. salinarum*. The halophile mother machine design used here is the same as those used previously for bacteria (Hussain et al., 2018), consisting of linear channels 1.5 μm wide, 1 μm deep, and a selection of lengths (80 μm used here; Supplementary Figure S4). The mold for the chip was fabricated via photolithography. The chip is arrayed with ~200 troughs per feeding channel (4 feeding channels per slide). Each chip was connected to microfluidics, supplying the growing cells with fresh medium throughout the course of each experiment (Methods). Under steady state growth conditions in the mother machine, Δ*ura3* cell area doubling time (6.85 ± 1.98 hr, Figure 4A) was similar to that in batch culture (6.68 hr; Figure 1B; Supplementary Table 3). Mother machine growth rates also reflected previous chamber growth measurements (6 ± 1 hr, (Eun et al., 2018)). Like the square chambers, the mother machine supports up to 6 generations of growth (Supplementary Movies 1-4). We conclude that the *Hbt. salinarum* parent strain grows optimally in the mother machine.

**Figure 4.**
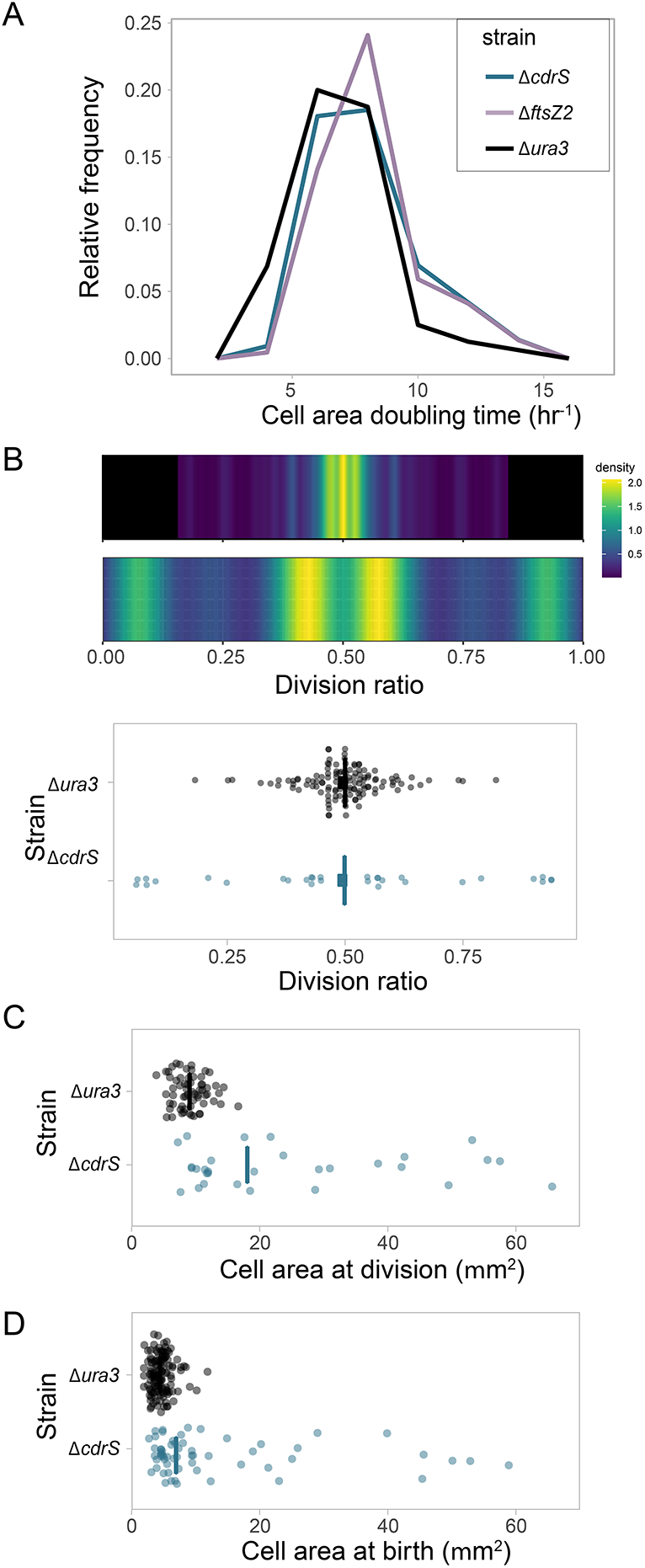
CdrS is required for division accuracy but not elongation. (A) Doubling time frequency plots for Δ*ura3* (black line), Δ*ftsZ2* (purple line), and Δ*cdrS* (green line). Legend colors are consistent throughout the figure. (B) Heatmap depicting the density distributions of division ratios for Δ*ura3* (top) and Δ*cdrS* (bottom). Cool colors represent low density, hot colors high density (see scale at right). Raw data dot plot shown below. Vertical crossbar represents the geometric mean. (C) Dot plot depicting area of mother cells at the time of division. Crossbar represents the median. (C) Dot plot depicting area of daughter cells immediately following division. Raw data dot plot shown below. Crossbar represents the median.

To compare division events from single cells across strains in real time using the mother machine, phase contrast time lapse images were used to quantify the cell growth parameters of elongation rate, division ratio, and interdivision time. Cell area doubling time (proportional to elongation rate) of the Δ*cdrS* (geometric mean 7.88 hr ± 2.21 hr) and Δ*ftsZ2* (8.04 ± 2.03 hr) mutants was statistically indistinguishable from the Δ*ura3* parent strain (6.86 ± 1.98 hr; Figure 4A; Supplementary Figure S5; *p*-values comparing each mutant to the parent > 0.76; effect size 0-0.1). This corroborates batch culture results that CdrS and FtsZ2 are not required for cell elongation. In contrast, division was strongly impaired for each mutant relative to the parent strain. Division of the Δ*cdrS* strain (30 of 108 cells divided) was observed at 35% the frequency of Δ*ura3* (63 of 80 cells), and the Δ*ftsZ2* strain was not observed to divide (0 of 110 cells). Δ*cdrS* cells that did not divide continued to elongate throughout the experiment, filling the chamber. In some cases, growth continued after the cell pole was extruded outside the single cell trough (Supplementary Movie SM1). Typically, Δ*ura3* cells divided in the center, with the division ratio (area daughter : area mother) centered at 0.492, variance CV = 17.6%, quantitatively consistent with previous observations (Eun et al., 2018). In contrast, Δ*cdrS* cells divided asymmetrically, with few cells observed to divide in the center (Figure 4B). This asymmetric division pattern was not random: 30% divided nearby the cell pole (division ratio <= 0.20), 50% divided offset from the cell midline (i.e. division ratio 0.3-0.6), a few outliers at the cell quarters (~0.25 or 0.75), and none dividing within 0.05 μm of the center (Figure 4B). The mean area of the Δ*cdrS* mother cells at the time of division (geometric mean 19.64 μm^2^; σ = 17.57) was significantly larger and more variable than that of Δ*ura3* (Figure 4C; 8.79 μm^2^; σ = 2.52; Welch’s *p*-value 1.7 × 10^−7^, effect size 1.50). Δ*cdrS* daughter cell mean area was twice as large as that of Δ*ura3* (Figure 4D; 8.81 μm^2^, σ = 13.67 vs 4.32 μm^2^, σ = 1.60, respectively, Welch’s p-value 2.3 × 10^−10^ and effect size 1.16). This suggests that asymmetric division leads to variable cell sizes of mothers and daughters, with a tendency toward larger cell size in Δ*cdrS* relative to that of the Δ*ura3* parent (Figure 4C and D).

In the same cells visualized for quantitation with phase contrast imaging, we tracked FtsZ1 division rings to differentiate active division events from cell fragmentation. The gene encoding monomeric superfolder GFP (msfGFP) was integrated at the native chromosomal locus by translational fusion to FtsZ1 in each of the three strain backgrounds (Methods). In the Δ*ura3* strain, we observed that cell division was preceded by helical assembly of the FtsZ1 ring (Figure 5A, Supplementary movie SM2). In contrast, deletion of *ftsZ2* abrogated ring formation in some cells, with a diffuse msfGFP-FtsZ1 signal observed throughout. In other Δ*ftsZ2* cells, rings formed but constriction was not observed (Supplementary movie SM3). Despite these division defects, Δ*ftsZ2* cells continued to elongate for the duration of the imaging experiments, filling the chamber (SM3). For Δ*cdrS* cells that were able to divide (though at a lower frequency than Δ*ura3* cells), each division event was preceded by formation of a msfGFP-FtsZ1 ring (Figure 5B; Supplementary Movie SM4a). However, not all ring formation resulted in division: in Δ*cdrS* cells that were not observed to divide, rings often formed but later disassembled (Figure 5C; Supplementary Movie SM4b). Taking these fluorescence images together with the quantitative analyses of the phase contrast images (Figure 4), these data strongly suggest that CdrS is an important regulator of cell division, and FtsZ2 is required for triggering cytokinesis at mid-cell. However, elongation appears to proceed independently of these factors.

**Figure 5.**
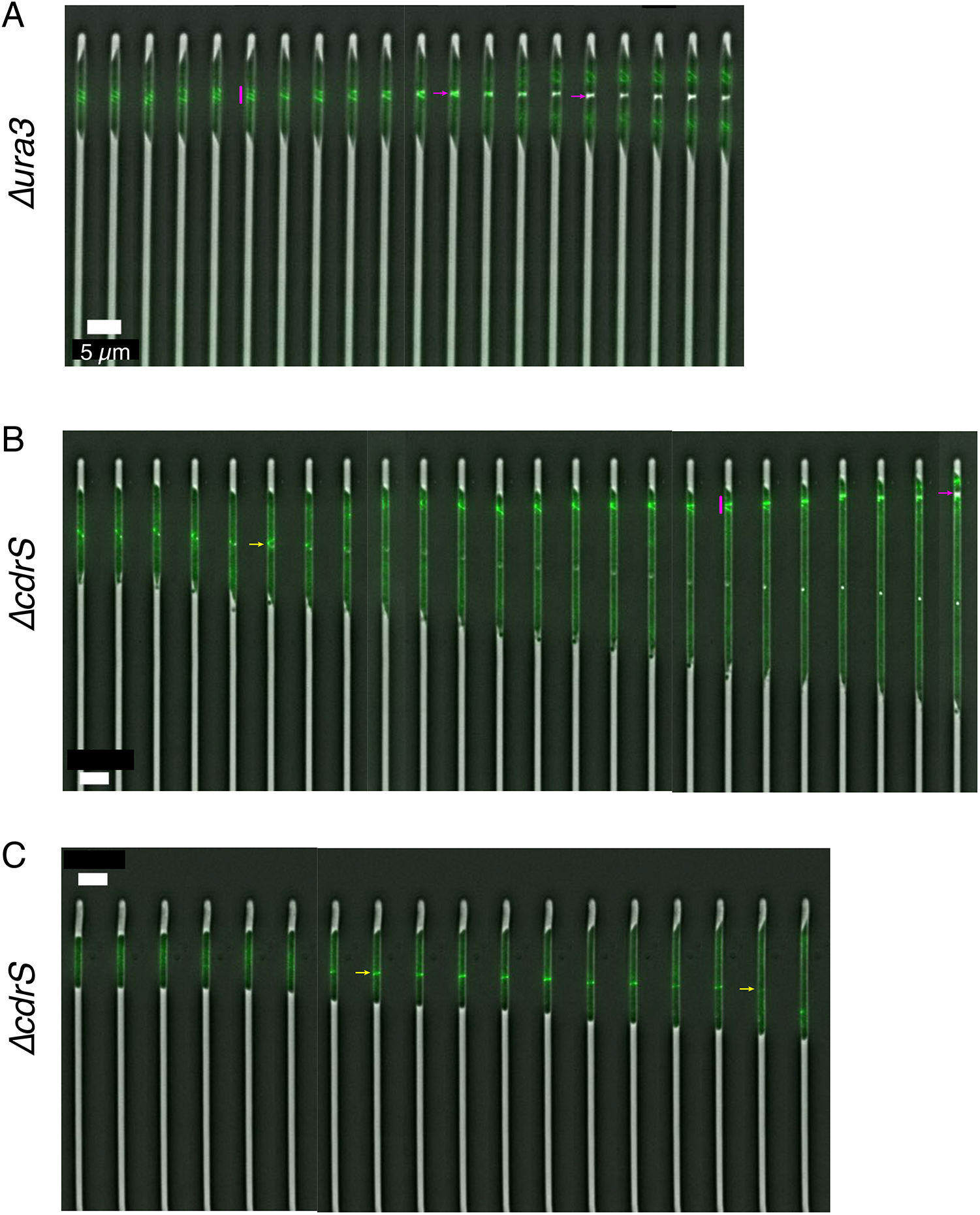
CdrS is important for triggering cytokinesis. (A) Montage of a representative Δ*ura3* cell growing in the mother machine. Pink vertical bar indicates the helical assembly of the division ring, and pink arrows indicate the division site just prior and just after division. Montage corresponds with Supplementary Movie SM2. (B) Montage of a representative Δ*cdrS* cell growing and undergoing polar division. Yellow arrow represents a division ring forming that later dissipates and does not result in division. Pink bar and arrow are as in panel A. Montage corresponds with Supplementary Movie SM4a. (C) Montage of a representative Δ*cdrS* cell that does not divide in the time frame of the movie. Yellow arrows represent a division ring forming and later dissipating, respectively. Montage corresponds with Supplementary Movie SM4b. For all panels, time between each still image is 20 minutes and scale bars are 5 μm.

### CdrS specifically regulates cell division and other cell cycle genes

To determine how CdrS regulates cell division, we compared gene expression of the Δ*cdrS* strain to the isogenic parent Δ*ura3* over the growth curve and in response to cell cycle arrest by aphidicolin. Given the cell division defects of the Δ*cdrS* strain, we focused on 20 genes known or predicted to be involved in growth and division in other systems, including all known *ftsZ* and *cetZ* paralogs encoded in the *Hbt. salinarum* genome. We used NanoString probe-based mRNA counting technology. Previous work demonstrated that this method is successful for accurate quantification of gene expression over time in *Hbt. salinarum* (Todor et al., 2013). Of the genes tested, 10 were significantly differentially expressed in the Δ*ura3* parent strain in batch cultures over the course of the growth curve, including *ftsZ2* and three CetZ homologs: *cetZ1 (VNG1933G), cetZ2 (VNG0265G) and cetZ5 (VNG6260G)* (Supplementary Figure S6A; Supplementary Table 4). Relative to the Δ*ura3* control strain, *ftsZ2, cetZ1,* and *sojA* were significantly differentially expressed in response to *cdrS* deletion during growth (Figure 6A). The protein product of plasmid-encoded *sojA* is a predicted member of the SIMIBI superfamily (NCBI accession cl28913), encompassing NTP-ases involved in a wide array of cellular functions, including the plasmid partitioning ParA AAA-type ATPase widely conserved in bacteria (Gerdes 2010 review). *ftsZ1* was not differentially expressed either over the growth curve or in the Δ*cdrS* vs the parent strain (Figure 6A, upper left). In contrast, *ftsZ2*, *cetZ1*, and *sojA* were dynamically expressed throughout growth, and expression levels were lower in Δ*cdrS* during early log phase (Figure 6A). For example, *ftsZ2* expression steadily increased ~1.8-fold throughout growth, reaching its peak during the transition to stationary phase (Figure 6A, upper right; Supplementary Table 4). In the Δ*cdrS* strain, *ftsZ2* expression followed a similar growth-dependent expression pattern. However, expression magnitude ranged from 1.3 to 2.2-fold lower across all growth time points in the Δ*cdrS* strain, with the largest defect in gene activation observed at the early log time point, with *cetZ1* and *sojA* following similar patterns (Figure 6A).

**Figure 6.**
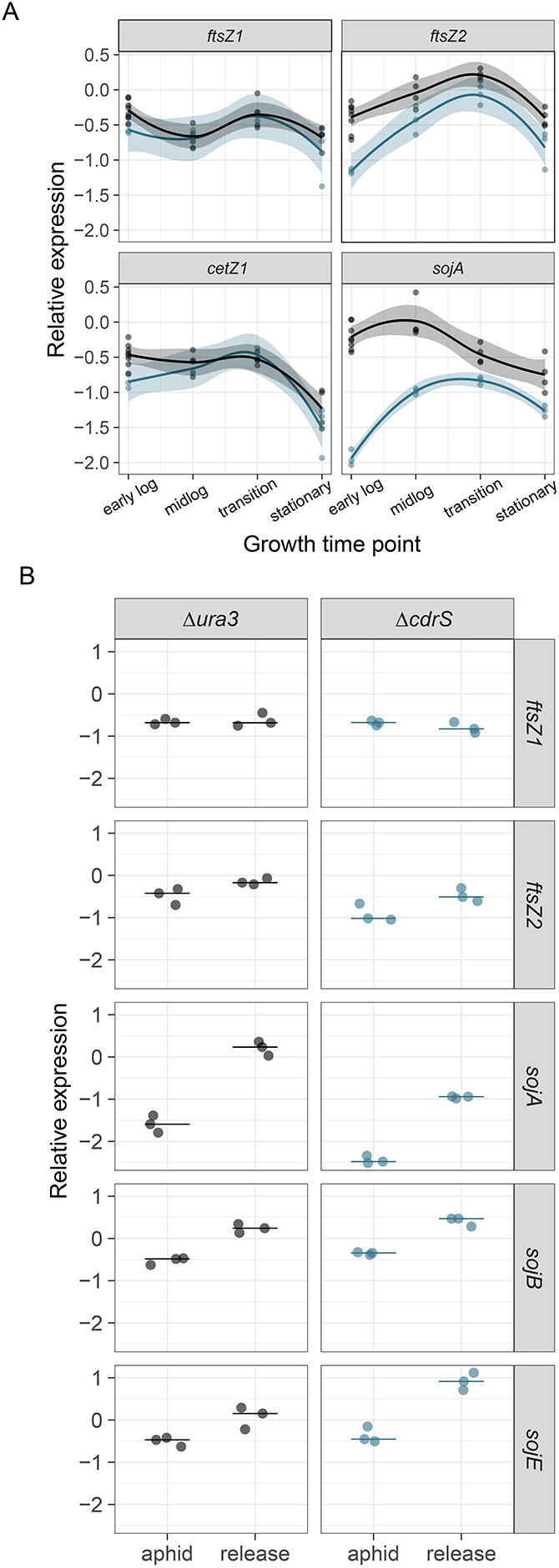
CdrS is important for wild type expression levels of growth and cell division genes. (A) Nanostring gene expression over the growth curve. Points represent log10 normalized expression data. Shaded regions represent smoothed conditional means. (B) Expression data in response to cell division block with aphidicolin. Labels on X-axis: aphid, gene expression during aphidocolin block; release, expression following wash-out. Points represent raw data, lines represent median.

In response to cell division block by aphidicolin and subsequent release into growth, 11 genes were significantly differentially expressed in the Δ*ura3* parent strain (Supplementary Figure S6B, Supplementary Table 4). *ftsZ2* was also significantly reduced in expression in the Δ*cdrS* strain relative to the Δ*ura3* strain in response to aphidicolin (Figure 6B). Three *par* family paralogs (*sojA, B,* and *E*) were also significantly mis-regulated in the Δ*cdrS* strain relative to the parent control under these conditions (Figure 6B). Together these expression data indicate that CdrS is important for wild type expression magnitude but not growth-dependent expression change of *ftsZ2, cetZ1,* and SIMIBI family protein-coding genes. These results are consistent with the hypothesis that CdrS is a specific regulator of the cell division ring and other putative cell division-related functions.

### CdrL is a specific and direct regulator of the *cdrS-ftsZ2* operon

To further investigate how the *cdrS-ftsZ2* locus is regulated, we conducted protein-DNA binding analysis by chromatin immunoprecipitation coupled to sequencing (ChIP-seq, see Methods). The putative DNA binding protein CdrL is encoded directly upstream of the *cdrS-ftsZ2* operon (Figure 1). Given this synteny and *cdrL* conditional co-expression with *cdrS-ftsZ2*, we reasoned that CdrL may play a role in regulation of the locus. The FLAG epitope was integrated into the chromosome at the 3’ end of the native *cdrL* locus, with the resultant strain encoding a C-terminal CdrL-FLAG translational fusion (Methods, Supplementary Table 5). In both mid-logarithmic and stationary phases of growth, the region upstream of the *cdrS-ftsZ2* locus was the only significant CdrL binding site reproducibly detected throughout the entire genome (Figure 7). Significant binding at other locations was detected in some ChIP-seq samples. However, these binding events were detected in redundant genomic regions, poor coverage in the input sample, or not detected across replicate samples (Supplementary Figure S7). We conclude that CdrL is a specific and direct regulator of *cdrS-ftsZ2* expression, binding exclusively and reproducibly upstream of this locus.

**Figure 7.**
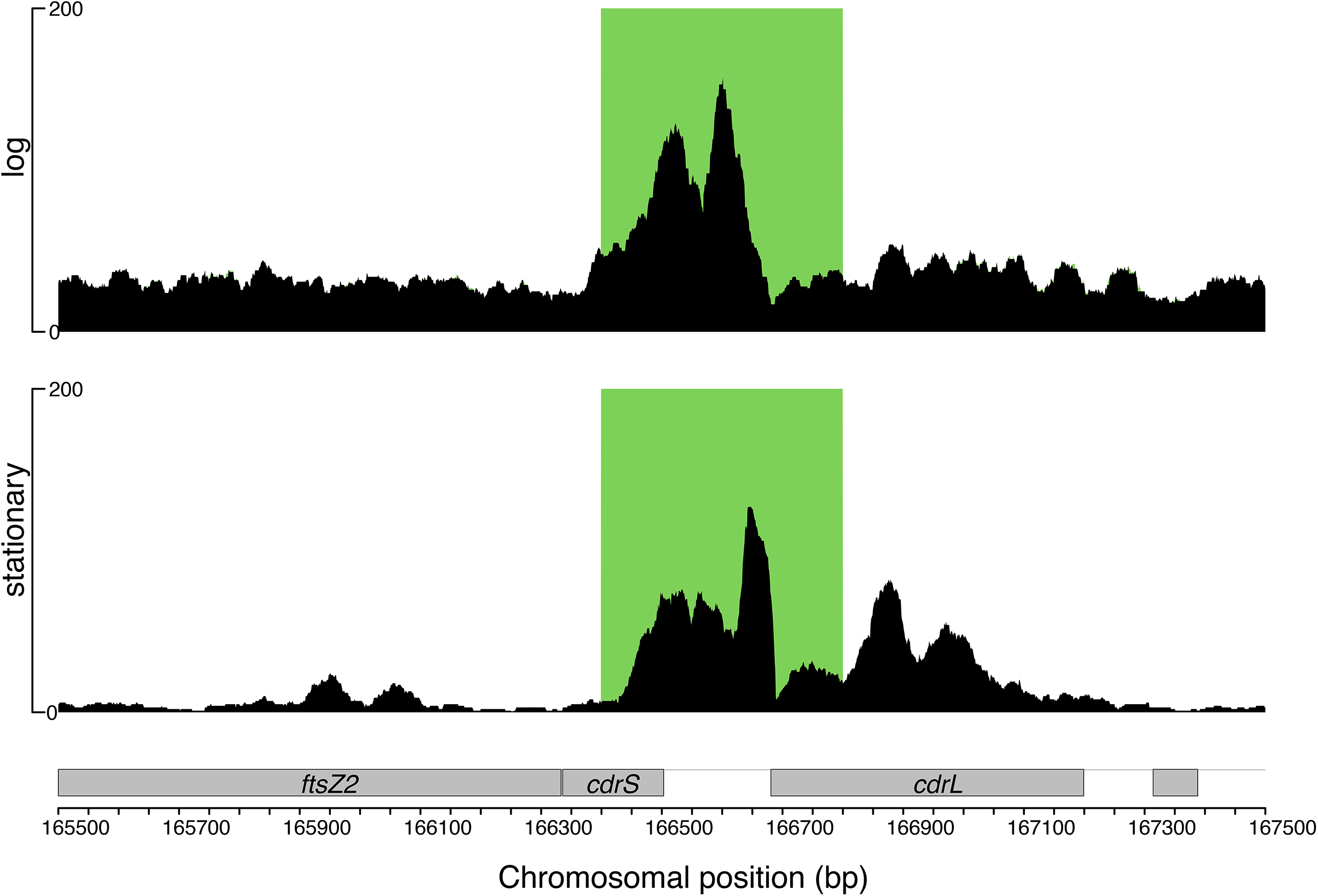
CdrL binds to the promoter region upstream of the *cdrS-ftsZ2* operon. Raw sequencing data for immunoprecipitated samples are shown by the black traces. Overlaid green boxes represent genomic regions detected by the peak detection algorithm (see Methods) to be significantly enriched for binding of CdrL relative to the input control. Y-axis scale represents read counts. CdrL binding site is shown for logarithmic phase cells (top) and stationary phase (bottom). Grey labeled boxes at bottom represent genes (reverse strand).

### *cdrS* homologs in other Haloarchaea are required for maintaining cell shape and size

The *cdrS-ftsZ2* locus was detected in all known Haloarchaeal genomes (Figure 1), and protein alignments showed strong conservation in model species across the clade (Figure 8A). The beta-sheet region of the RHH protein was perfectly conserved, and only 9 residues of the alpha helical regions varied across these species (Figure 8A). Given this strong conservation, we hypothesized that CdrS plays a conserved functional role as a regulator of cell division across hypersaline-adapted archaeal species. However, multiple attempts to delete *cdrS* in the genetically amenable model species *Haloferax volcanii* [HVO_0582; HVO_RS07500] and *Haloferax mediterranei* [HFX_0561, HFX_RS02725] were unsuccessful despite using a selection-counterselection scheme routinely used in the field (Allers et al., 2004; Liu et al., 2011). In *Hfx. volcanii,* 48 clones were screened by PCR across 3 transformations, and 292 clones across 4 transformations in *Hfx. mediterranei*. No *Hfx. mediterranei* knockout candidate clones were detected. Eight *Hfx. volcanii* candidates were identified; however, Sanger sequencing detected many point mutations throughout the locus. These results, corroborated by a parallel study on *cdrS* in *Hfx. volcanii* (Vogel et al, 2020), suggest that *cdrS* is required for viability under laboratory conditions in these species.

**Figure 8.**
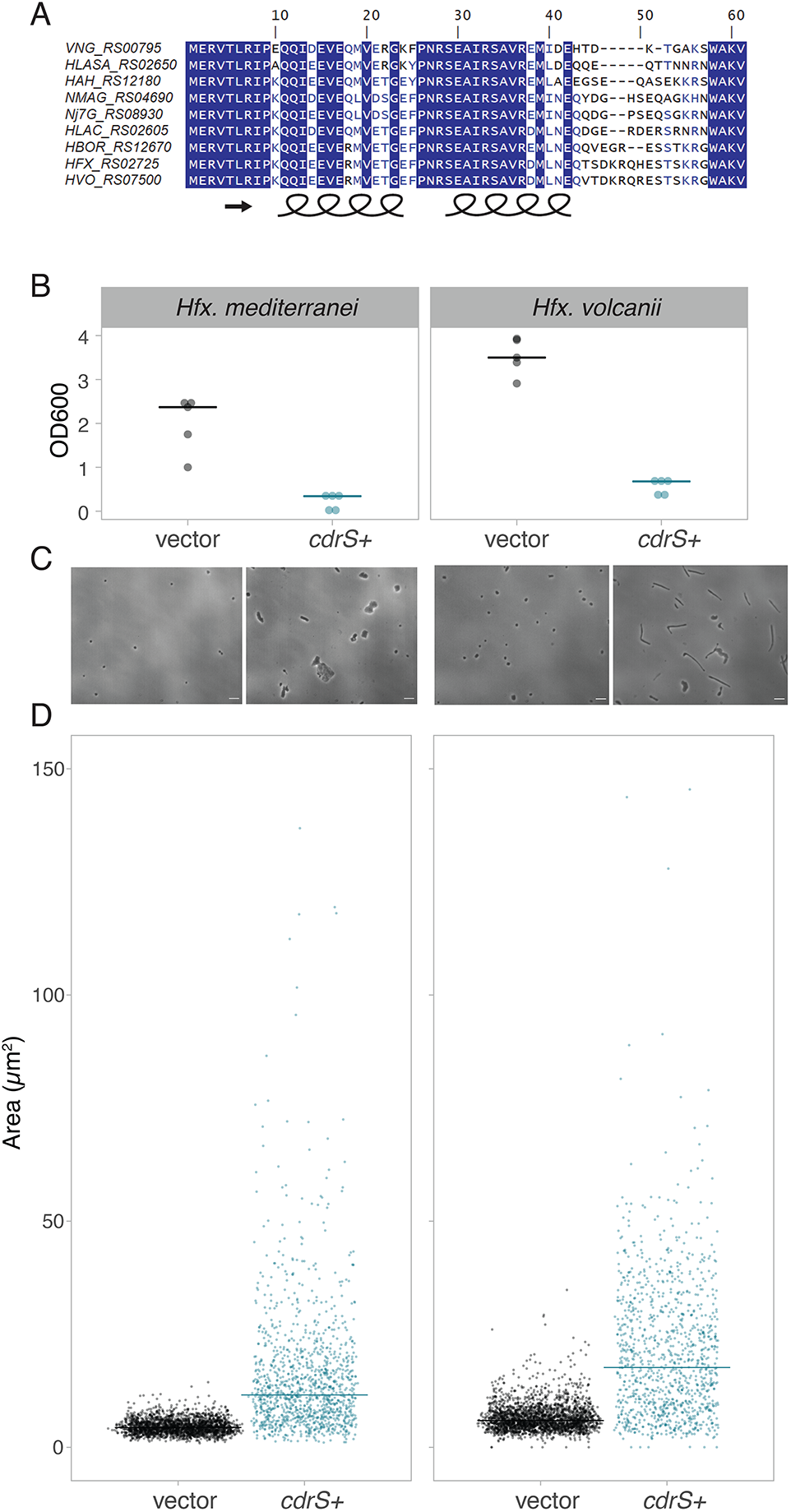
CdrS is required for cell division across halophiles. (A) Clustal Omega alignment of protein sequences from halophilic archaeal model organisms. The GenBank protein sequence identifiers for CdrS homologs are given at left. Species identifiers are as follows: VNG, *Hbt. salinarum*; HLASA, *Halanaeroarchaeum sulfurireducens*; HAH, *Haloarcula hispanica*; NMAG, *Natrialba magadii*; Nj7G, *Natrinema sp.* J7-2; HLAC, *Halorubrum lacusprofundi*; HBOR, *Halogeometricum borinquense*; HFX, *Hfx. mediterranei*; HVO, *Hfx. volcanii*. (B) Final cell density measurements of overnight cultures of empty vector control strains (pJAM202c, black) vs *cdrS*+ overexpression strains (blue) in *Hfx. mediterranei* (left, *HFX_0561*, pAKS77) and *Hfx. volcanii* (right, *HVO_0582*, pAKS78). (C) Phase contrast micrographs comparing empty vector control to *cdrS*+ overexpression strains. Species and strain labels align in columns according to labels in panel B. (D) Quantification of cell area comparing the empty vector control strain (vector, black dots) to *cdrS* overexpression strain (*cdrS*+, blue dots) in *Hfx. mediterranei* (left panel) and *Hfx. volcanii* (right panel). Three large *cdrS+* outlier cells in *Hfx. mediterranei* (area 218.44, 209.22, 221.86 μm^2^) are not shown for figure clarity. Crossbar indicates the median cell area.

Instead, we overexpressed *cdrS* in these species to investigate how its role in cell division is conserved. Both *cdrS*^*Hv*^ and *cdrS*^*Hm*^ were cloned downstream of the strong constitutive *Hbt. salinarum* rRNA P2 promoter in vector pJAM202c and transformed into the respective *Haloferax* species (Kaczowka and Maupin-Furlow, 2003). Overnight cultures of the empty vector control strain grew well under selection, reaching high final cell densities (OD600 2.0-3.5; Figure 8B). In contrast, both *Haloferax* species carrying the *cdrS* overexpression plasmid (*cdrS+*) exhibited significant growth inhibition (Figure 8B), indicating the importance of tight control of *cdrS* expression levels. In both species, severe morphological defects were observed in *cdrS*-overexpression strains compared to the disc-shaped control strain (Figure 8C). *Hfx. volcanii cdrS+* overexpression cell area was, on average, 3-fold larger than the corresponding empty vector control strain (Welch’s *p* < 2.2 × 10^−16^; effect size 1.355 (large); Figure 8D; Table 3). *cdrS*+ cells were also 3-fold longer than the disc-shaped control, suggesting that the area increase was primarily due to elongation of the cell body. However, thickness varied along the length of the *cdrS+* cells, often resulting in club-like shapes (Figure 8C). Similar to *Hbt. salinarum* Δ*cdrS* (Figures 2, 4), *cdrS+* cell area was more variable than that of the empty vector cells, suggesting impaired regulation (Table 3, Figure 8D). Similar results were obtained with the *Hfx. mediterranei* system, with 3-fold larger cell area and increased variance observed relative to the control strain (Welch’s *p* < 2.2 × 10^−16^; effect size 1.536 (large); Table 3; Figure 8C, D). However, unlike cdrS^*Hv+*^ overexpression cells, the *cdrS*^*Hm+*^ cells were, on average, only 1.5-fold longer than control cells. We observed that *cdrS*+ in *Hfx. mediterranei* exhibited two major forms, one with an increase in cell area across two planes, generating plate-like cells, and the other elongated and/or club-shaped (Figure 8C). Taken together, these data suggest that CdrS is required for maintaining wild type disc-like cell shape and size in other halophilic archaea. Given the similarity of these phenotypes with those of *Hbt. salinarum*, these data are consistent with the hypothesis that CdrS^*Hv*^ and CdrS^*Hm*^ are also required for cell division regulation but may play an additional role in maintaining cell shape and viability.

**Table 3.** Summary statistics of cell area and length in *Haloferax* species

|  |  | Area |  |  |  |
| --- | --- | --- | --- | --- | --- |
| Species | Strain | geometric mean | median | $\sigma$ | n |
| *Hfx. volcanii* | *cdrS*+ | 15.71926845 | 17.785675 | 49.54274 | 1105 |
| *Hfx. volcanii* | vector | 5.903139311 | 5.925052 | 2.964239 | 2554 |
| *Hfx. mediterranei* | *cdrS*+ | 11.64842328 | 11.634059 | 17.05703 | 1253 |
| *Hfx. mediterranei* | vector | 4.298809837 | 4.3827778 | 1.600874 | 2291 |
|  |  | Length |  |  |  |
| | | geometric mean | median | $\sigma$ | n |
| *Hfx. volcanii* | *cdrS*+ | 9.031154158 | 10.20501 | 8.200973 | 1105 |
| *Hfx. volcanii* | vector | 3.25569573 | 3.1692971 | 1.30773 | 2554 |
| *Hfx. mediterranei* | *cdrS*+ | 5.32907322 | 5.2788867 | 3.555722 | 1253 |
| *Hfx. mediterranei* | vector | 2.619884171 | 2.6185364 | 0.547168 | 2291 |

## Discussion

Growth and division are precisely controlled to ensure the coordination of cellular events. However, such regulation thus far has been unexplored in archaea. In the current study, we identify and characterize the highly conserved CdrSL gene regulatory network (GRN). Quantitative microscopy on cells from bulk culture and single cell time lapse images demonstrates the requirement of CdrS for cell division in the model archaeal species *Hbt. salinarum* (Figures 2-5). Specifically, deletion of *cdrS* or *ftsZ2* impairs cell division but does not affect cell elongation rate, providing strong evidence that the CdrS regulatory system and FtsZ2 itself are required for coupling cell growth and division. Intriguingly, while FtsZ2 is absolutely required for cell division, a small subset of Δ*cdrS* cells are able to divide (Figures 4, 5), hinting that other mechanisms regulating archaeal cell division await discovery.

Our genetic evidence, transcript profiling, and protein-DNA binding suggest that regulation is achieved via CdrS transcriptional activation of genes that encode proteins predicted to function in critical aspects of cell division (Figures 6, 7). These include the cell division ring (FtsZ2), cell shape maintenance (CetZ1; (Duggin et al., 2015)), and DNA partitioning (SojABE). One caveat is that the *soj* genes are encoded on the *Hbt. salinarum* pNRC100 and pNRC200 megaplasmids. These genomic elements are subject to frequent copy number variation (Dulmage et al., 2018), so further evidence is needed to determine the definitive mechanism by which CdrS affects their expression. Nevertheless, CdrS exerts its effect on these genes in early log phase and following release from a chemical cell division block, suggesting that CdrS acts during the transition from stasis to growth. CdrL provides a second level of regulation by binding the region upstream of the *cdrS-ftsZ2* operon (Figure 7). Together these data suggest a mechanism by which the CdrSL system controls cell division.

Here we show that FtsZ proteins have distinct but interrelated roles in cell division, which are reflected in their differential regulation by CdrS (Figures 4-6). Previous work in halophilic archaea suggest that a large class of tubulin-like proteins, FtsZ and CetZ, function in cell division and cell morphology, respectively (Aylett and Duggin, 2017; Duggin et al., 2015). Here we build on this knowledge, demonstrating that *ftsZ1* expression levels are independent of CdrS regulation, and remain fairly constant at different growth rates (Figure 6). FtsZ1 rings form just prior to cell division events (Figure 5). In contrast, *ftsZ2* transcript levels fluctuate depending on the presence of CdrS, growth phase, and chemical perturbations, indicating a growth-sensitive mechanism of transcriptional regulation (Figure 6). Although *ftsZ2* can be deleted in *Hbt. salinarum* (Figures 2-5, Supplementary Table 2), it is essential for triggering the constriction of the cytokinetic ring during exponential growth, further suggesting condition-specific functions for FtsZ2 (Figure 5). Therefore, the multiple copies of FtsZ within the Halobacteria, and likely other Euryarchaeota, do not appear to be redundant but instead may represent a case of subfunctionalization. CdrS plays a key role in regulating the interrelated but separate functions of the two FtsZ proteins, as Δ*cdrS* cells fail to activate *ftsZ2* during rapid growth (early log phase), delaying cell division (Figures 4-6). Given the independence of cell elongation from regulation by CdrS and FtsZ2 (Figures 2-4), and that haloarchaeal cells grow by inserting new surface-layer (S-layer) material at midcell (Abdul-Halim et al., 2020); cell elongation and cytokinesis may occur at the same cell region (Z ring), but may be temporally sequential and separately regulated. CdrS appears to play a key role in coordinating these events.

Our results evoke a function for archaeal FtsZ proteins analogous with those required for chloroplast division in land plants and some bacterial species. In chloroplasts, FtsZ1 and FtsZ2 have interrelated but non-redundant functions in cell division. The two FtsZ homologs form co-polymers, with one thought to be involved in divisome structure, the other involved in dynamic GTP turnover and constriction (TerBush et al., 2013). Similarly, in the alpha-proteobacterium *Agrobacterium tumifaciens*, although two FtsZ proteins co-polymerize at mid-cell, only one is required for constriction (Howell et al., 2019). Recent results in *Hfx. volcanii* suggest co-localization of FtsZ1 and FtsZ2 at mid-cell (Liao et al., 2020). Consistent with these multi-FtsZ models of cell division, the defects observed here in *Hbt. salinarum* FtsZ1 ring assembly in the absence of FtsZ2 or CdrS suggest that FtsZ1 and FtsZ2 could also form co-polymers whose stoichiometry is balanced by CdrS regulation.

The CdrS-FtsZ2 system is widely conserved across the Archaea at both the protein structural and functional levels, with CdrL restricted to the Halobacteria (Figures 1,8). At the level of primary structure, the ribbon-helix-helix (RHH) CdrS protein is detected in every major known taxonomic group of archaea except DPANN (Figure 1). Further, RHH protein CdrL is encoded in all sequenced members of Halobacteria, though often only annotated by the C-terminal double zinc ribbon (DZR) domain. The CdrS and CdrL clades are phylogenetically distinct within the RHH superfamily (PF01402, Figure 1, Supplementary Figure S1), suggesting independent evolutionary history. Restriction of *cdrL* to the Halobacteria further supports this hypothesis. Therefore, we predict that the locus acquired *cdrL* after the divergence of Methanomicrobial and Halobacterial ancestors.

CdrS is also conserved at the functional level, as we demonstrate that CdrS is required for proper cell division across multiple species of haloarchaea, including *Hbt. salinarum*, *Hfx. volcanii*, and *Hfx. mediterranei* (Figure 8). This was corroborated independently in a companion study, which also demonstrated that *cdrS* is an essential gene whose product is required for cell division in *Hfx. volcanii* (Vogel et al., 2020). Given the essentiality of *cdrS* in both *Haloferax* species and the polyploidy of Halobacteria, we have included whole genome sequencing (WGS) as an essential step of strain construction (Supplementary Table 2). Particularly with seemingly essential genes, we have found WGS more sensitive than standard PCR and Sanger sequencing in detecting all copies of a target gene as well as ruling out second site suppressor mutations (see also (Zaretsky et al., 2019)). Taking our phylogenetic and genomics evidence together, we conclude that the CdrS-FtsZ2 system is widely conserved and important for cell division across hypersaline adapted archaea.

## Methods

### Bioinformatic prediction and phylogenetic analysis

Protein structural predictions of CdrS and CdrL were conducted using the Phyre2 server (Kelley et al., 2015) using default parameters, access date 2/12/2020. The top hit reported in the text was the structure CopG DNA binding domain of the NikR transcription factor [protein databank identifier 2BJ7 [PDB, rscb.org, access date 2/12/2020, original structure published in (Chivers and Tahirov, 2005)]. Protein primary sequence predictions are reported for protein families [PFAM; pfam.xfam.org; (El-Gebali et al., 2019); access date 2/12/2020], e-values of significance of the matches for each protein were found in the *Hbt. salinarum* genome database ((Ng et al., 2000); https://baliga.systemsbiology.net/projects/halobacterium-species-nrc-1-genome/). Gene expression correlations were calculated and visualized (Figure 1C) using the corrplot and psychometric packages in the RStudio coding environment, R version 3.6.1. Synteny of the *cdrL-cdrS-ftsZ2* ((Ng et al., 2000) identifiers *VNG0195H-VNG0194H-VNG0192G*; NCBI identifiers *VNG_RS00800-VNG_RS00795-VNG_RS00790*) locus was determined using the SyntTax database using default parameters [https://archaea.i2bc.paris-saclay.fr/SyntTax/; accessed January 2019; (Oberto, 2013)]. All 384 archaeal genomes housed in the SyntTax database were searched with the FtsZ2 (VNG0192G) protein sequence of *Hbt. salinarum*. Detection of locus homologs and synteny for those genomes not included in SyntTax (Bathyarchaeota, Korarchaeota, Asgard, 20 genomes) were found using NCBI genomes database using BLAST to detect FtsZ2 homologs (sequence similarity >200 bits). Subsequent manual inspection in the NCBI genome browser (https://www.ncbi.nlm.nih.gov/genome/) detected synteny of the locus. Locus identifiers for *cdrS-ftsZ2* across 93 archaeal genomes; and UniProt protein identifiers for FtsZ-family homologs (including FtsZ and CetZ-like proteins) in the absence of *cdrS* across 1,497 genomes are given in Supplementary Table 1. CdrS protein sequence alignments shown in Figure 8 were conducted using Clustal Omega using default parameters in the DNAstar MegAlign software package.

### Strains, plasmids, and primers

*Halobacterium salinarum* NRC-1 (ATCC 700922) was the wild-type used in this study. Gene deletions and chromosomal integrations were performed using two-stage selection and counterselection homologous recombination in the Δ*pyrF* (Δ*ura3*) strain isogenic parent background as described (Peck et al., 2000), updated in (Wilbanks et al., 2012), and subject to whole genome resequencing here (Supplementary Table 2, Sequence Read Archive PRJNA614648). Plasmids were constructed using isothermal assembly (Gibson, 2011) and propagated in *E. coli* NEB5α; see primer list for details (Supplementary Table 6). Primers were ordered from Integrated DNA Technologies (Coralville, IA). Final strain genotypes were verified using site-specific PCR and Sanger sequencing by Eton Biosciences, Inc (San Diego, CA) and genomic DNA extraction followed by Illumina sequencing (see section below). Plasmids used in cloning are presented in Supplementary Table 7 and resultant strains in Supplementary Table 5.

For overexpression studies, *Haloferax volcanii* DS2 and *Haloferax mediterranei* ATCC33500 were the wild-type strains. Plasmids were constructed from pJAM809 using restriction enzymes NdeI and KpnI to remove the resident ORF and replace with the *cdrS* gene from each species. Δ*pyrE2* derivatives of *Haloferax* species (Bitan-Banin et al., 2003; Liu et al., 2011) were transformed with NEB5α-propagated plasmid. Due to concerns about the higher mutation rate in methylase-deficient *E. coli*, we opted not to passage plasmids through a *dam*^−^/*dcm*^−^ strain as is commonly done (Dyall-Smith, 2009).

### Media and growth conditions

*Hbt. salinarum* strains were routinely grown using CM medium (250 g/L NaCl (Fisher Scientific); 20 g/L MgSO_4_·7H_2_O (Fisher Scientific); 3 g/L trisodium citrate (Fisher Scientific); 2 g/L KCl (Fisher Scientific); 10 g/L Bacteriological Peptone (Oxoid); pH 6.8)). Media were supplemented with 50 μg/ml uracil (Sigma) to complement the uracil auxotropy of the Δ*ura3* background. During knockout and integrant strain construction, the first stage of selection was performed on mevinolin (10 μg/ml; AG Scientific) plates (CM with 20 g/L agar; Difco), and the second stage of counterselection on 5-Fluoroorotic acid (300 μg/ml; ChemImpex) were used in agar plates. All growth was performed at 42°C and liquid culturing was shaken at 225 rpm under ambient light. Self-replicating plasmids were maintained using 1 μg/ml mevinolin in liquid culture. *E. coli* was grown in LB with carbenicillin (50 μg/ml; Sigma) to maintain plasmids. Maximum instantaneous growth rates were calculated as described in (Sharma et al., 2012). Raw data are provided in Supplementary Table 3.

For statistical analysis of OD vs CFU data shown in Figure 2, we noted that the slopes of the regression lines were similar between Δ*ura3* and each mutant, so we fit a linear model using log_2_(OD) and genotype to predict log_10_(CFU). In a two-way ANOVA test, we found no evidence of interaction between strain and OD vs CFU slope (Δ*ura3* vs Δ*cdrS*: *p*-value = 0.225; Δ*ura3* vs Δ*ftsZ2*: *p*-value = 0.95). Therefore, a new model was fit that constrained equal slopes for the regression line for each strain, allowing us to determine the difference in CFU/ml per unit log_2_(OD) between strains. These differences are reported in the text.

### gDNA extraction and Illumina sequencing

*Hbt. salinarum* strains were grown to mid-logarithmic phase (OD_600_ ~0.7) and 1 mL pelleted by centrifugation. Pellets were stored at −20°C until processed. DNA was extracted using a phenol-chloroform method. Briefly, pellets were lysed in dH_2_O and treated with RNase A and Proteinase K. DNA was extracted using phenol:chloroform:isoamyl alcohol (25:24:1) (Fisher Scientific) in Phase Lock Gel tubes (QuantaBio) and ethanol precipitated. DNA pellet was resuspended in modified TE buffer (10 mM Tris-HCl pH 8.0, 0.1 mM EDTA). Purified DNA was quantified using a Nanodrop systems (Thermo Scientific) and sonicated in a Diagenode sonicating water bath for 20 cycles on high. DNA quality was assessed by Bioanalyzer using a High Sensitivity DNA chip (Agilent). Samples were submitted to Duke Center for Genomic and Computational Biology core sequencing facility for adapter ligation with TruSeq (Illumina) adapters and library amplification. Samples were pooled and run in a single lane on an Illumina HiSeq 4000. 50 bp reads were assessed for quality using FastQC and adapter sequences trimmed using Trim Galore and Cutadapt. Reads were aligned to *H. salinarum* NRC-1 genome using Bowtie2 within the breseq package as described in (Deatherage and Barrick, 2014; Zaretsky et al., 2019). breseq results were analyzed for incomplete gene conversion, SNPs, and possible genomic rearrangements. Results are given in Supplementary Table 2, code freely available on github https://github.com/amyschmid/aglB-WGS-growth, and raw data accessible through the Sequence Read Archive (SRA) accession PRJNA614648.

### Microscopy and quantitative analysis of cell morphology

For microscopy of the deletion mutants, Δ*cdrS*, Δ*ftsZ2*, and Δ*ura3* cells were collected at various stages throughout the growth curve (see Supplementary Table 3 for OD600 at harvest) or in response to aphidicolin treatment (see subsection below). In each experiment, 5 μl each of 3-5 biological replicate culture samples were mounted on 1% agarose pads equilibrated with basal salt buffer (CM medium without peptone) overnight at room temperature. Phase contrast images were taken using a Zeiss Axio Scope.A1 microscope (Carl Zeiss, Oberkochen, Germany) and a PixeLINK CCD camera (PixeLINK, Ottawa, Canada) at 40X magnification. Quantification of cell length and area was calculated using the Fiji distribution (Schindelin et al., 2012) of ImageJ (Rueden et al., 2017) and plug-in MicrobeJ (Ducret et al., 2016). For batch culture experiments (overexpression and deletion mutants), the significance of the difference in cell size (length and/or area) between mutant vs parent strain cells in was calculated using Welch’s modified t-tests on order quantile normalized data, and effect size of these differences calculated using Cohen’s D test. P-values were adjusted for multiple hypothesis testing using the Benjamini-Hochberg correction.

#### Aphidicolin treatment

Cells were cultured to stationary phase, then sub-cultured in 20 mL of CM medium supplemented with uracil to OD_600_ ~0.01. Subcultures were incubated at 42°C with 225 rpm shaking until OD_600_ ~0.3, at which point 30 μM aphidicolin (30 mM stock in DMSO, Sigma) was added. After 6 hours incubation with drug, the remaining culture was washed twice by centrifugation (4500 × *g*, 6 minutes), resuspension in fresh CM, and incubation for 20 minutes at 42°C with 225 rpm shaking. Samples were harvested for microscopy (5 μl) prior to aphidicolin addition, prior to washing, and 11 hours after washing (release) from aphidicolin treatment. Rationale for time point selection was based on (Herrmann and Soppa, 2002). Mixed effects ANOVA with Welch’s post-hoc t-tests on ordinary quantile normalized data with Benjamini-Hochberg correction were used to determine statistical significance of the differences between strains and timepoints.

#### Microscopy of overexpression strains across species of halophiles

Five biological replicate overnight cultures *Hfx. volcanii* strain H26 harboring the pAKS78 plasmid and *Hfx. mediterranei* WR510 harboring the pAKS77 plasmid were grown in rich Hv-YPC medium. Significance of the difference between *cdrS*+ overexpression strain cells and empty vector control strain in each species was calculated as described above for knockout mutants and aphidocolin experiments.

### Single cell microfluidics time lapse microscopy

#### Strains and growth conditions

All strains analyzed in the mother machine experiments were streaked fresh from −80°C stocks and inoculated in 50 mL of CM supplemented with uracil in beveled flasks and grown at 42°C to OD_600_ 0.6-0.8 before loading into the microfluidic chip.

#### Microfluidic chip fabrication

The microfabrication of the master mold used to create the microfluidic chip used in our mother machine experiments was as previously described in (Norman et al., 2013) except that the second, wider layer in the cell chambers meant to enhance growth was not necessary to maintain the cell’s growth over the timescale of the current experiment. Specifically, to fabricate the microfluidic device, the features were molded into a piece of polydimethylsiloxane (PDMS) by pouring dimethyl siloxane monomer (Dow SYLGARD 184 Silicone Elastomer Base) mixed with a curing agent (Dow SYLGARD 184 Silicone Curing agent) in a 10:1 ratio on top of the master mold, followed by degassing under a vacuum, and curing the set-up overnight at 65°C. The solidified PDMS piece was then peeled from the master and cut into approximately 1.5 × 1.5 cm chips. Access holes for each of the feeding channels were punched using a 0.75 mm biopsy punch (WPI). The PDMS chip was then bonded to a KOH-cleaned 22 × 60 mm glass coverslip (VWR, no. 1.5) by oxygen plasma treatment at 200 mTorr of pressure and 30 Watts for 30 seconds in a PE-50 compact benchtop plasma cleaning system (Plasma Etch). Chips were baked at 65°C for at least 1 hour before use.

#### *Loading* Hbt. salinarum *cells into microfluidic devices*

CM medium supplemented with 1% bovine serum albumin (BSA) was manually injected into the microfluidic device using a 1 mL syringe and incubated for 1 hour. To avoid the crystallization of salt from the media, microcapillary pipet tips with CM media were left attached to the access holes of the chip. Prior to loading, cell cultures were filtered through a 40 μm cell strainer (EASYstrainer, Greiner Bio-One) to remove large salt crystals formed during culture growth. Cultures were then centrifuged at 3000 × *g* for 5 minutes and concentrated to a final volume of 100 μL. Cells were manually loaded into the main flow channel of the microfluidic mother machine device using a 1 mL syringe. To increase the number of cells in the mother machine wells, the chip was briefly centrifuged in a VWR Galaxy mini centrifuge. The microfluidic chip was then connected to automatic Harvard Apparatus syringe pumps by Tygon tubing (Saint-Gobain, ID 0.020 in) and blunt end dispense tips (Fisnar, 21 gauge, 1 inch). Fresh medium was continuously pumped at 1-2 μL/min at 37°C for 30 minutes to allow cells to further propagate within the mother machine wells. The system was subsequently moved to the microscope for data collection.

#### Microscopy

Cells in the mother machine were imaged in a Nikon Eclipse Ti microscope with a 6.5μm pixel CMOS Hamamatsu camera and a Nikon 100X NA 1.45 phase-contrast objective. Images were captured every 20 minutes for 1-2 days. Exposure times for both phase contrast and fluorescence were 200 milliseconds. Epi-illumination was provided by a fiber coupled Agilent launch for 488nm to image msfGFP-FtsZ1 in AKS137, AKS170 and AKS196.

#### Single-cell growth analysis from the mother machine

Using phase-contrast images; 80 Δ*ura3*, 108 Δ*cdrS*, 110 Δ*ftsZ2* cells were manually traced in Fiji image analysis software (Schindelin et al., 2012) to determine cell area through pixel counting. These measurements were used to determine the cell area doubling (elongation) rates by fitting an exponential curve to the single cell growth rate graphs of change of the cell area over time. *Hbt. salinarum* has previously been shown to grow exponentially through single cell analysis (Eun et al., 2018). The area at birth was calculated from images immediately following complete separation of daughter cells. The area during division was determined by adding the area of the two daughter cells immediately after division. The division site placement was determined by treating each pole separately, then calculating the ratio of each daughter cell to its corresponding parent. Single cell measurements shown in the figures were plotted using ggplot2 (Wickham, 2016) in the RStudio coding environment. Statistical analyses were conducted using the sjstats (Ludecke, 2020) and rstatix packages in RStudio. Significance of differences in doubling time between strains were modeled using 3-way ANOVA, with effect sizes calculated using an η^2^ test. Significance of the differences between mother or daughter cells in the parent and mutant strains were calculated using Welch’s modified t-tests on order quantile normalized data, and effect size of these differences calculated using Cohen’s D test. P-values were adjusted for multiple hypothesis testing using the Benjamini-Hochberg correction.

### Gene expression analysis with NanoString

To collect RNA across the growth curve, culture aliquots were collected from flask cultures at 4 phases of growth (early logarithmic, mid-logarithmic, early stationary phase, and late stationary phase. 50 ml of culture was sampled at low OD_600_ (~0.05), 2 ml at higher OD_600_ (~2.0). RNA was collected following 6 hours of aphidicolin treatment and 11 hours after release as described above. Culture samples were centrifuged at 21,000 × *g* for 2 minutes, supernatant removed, and immediately frozen in liquid nitrogen. Pellets were stored at −80°C overnight and RNA was purified using an Absolutely RNA Miniprep kit per manufacturer's instructions (Aglient, Santa Clara, CA). To verify the lack of DNA contamination, end point PCR was conducted for 30 cycles on 200 ng of RNA sample using primers given in Supplementary Table 6. RNA quality was determined using a Bionanalyzer and RNA Nano 6000 chip according to manufacturer’s instructions (Aglient, Santa Clara, CA).

Gene expression was quantified using NanoString detection and a custom probe Codeset (Supplemental table 5; (Geiss et al., 2008)). Probes were designed to target 23 genes predicted to encode proteins involved in cytoskeletal and growth functions. One hundred nanograms of RNA was hybridized and quantified using the nCounter instrument by the Duke Microbiome Shared Resource core facility. Counts were normalized using three housekeeping genes (*eif1a2* [*VNG_RS06805*], *coxA2* [*VNG_RS02595*], *VNG1065C* [*VNG_RS04150*]) and NanoString nSolver software, then further normalized to expression in the NRC-1 wild type control. Significance of this relative normalized differential expression between the parent and Δ*cdrS* strain was assessed using the maSigPro package (Conesa et al., 2006) in the R Bioconductor coding environment with default parameters except: Q-value 0.01; alfa 0.01; R-squared cutoff 0.7. We conducted Benjamini-Hochberg correction for multiple hypothesis testing in the context of the maSigPro package. All raw and normalized data and probe sequences are available in Supplementary Table 4. R code is given in the github repository associated with this study https://github.com/amyschmid/cdr.

### ChIP-seq experiment and analysis

Triplicate cultures of strains AKS113 (CdrL tagged at the C-terminus with the FLAG epitope, Supplementary Table 5) and Δ*ura3* control strain were grown until stationary phase and subcultured in rich media supplemented with uracil. At mid-log phase (OD_600_ ~0.15) and early stationary phase (OD_600_ ~1.8), cultures were crosslinked and immunoprecipitated as described (Wilbanks et al., 2012) with the following exceptions: cultures were crosslinked with 1% formaldehyde for 30 minutes at room temperature; immuoprecipitations were conducted using Dynabead magnetic beads (Thermo-Fisher product 10002D) conjugated with anti-FLAG (Abcam ab1162) anti-rabbit monoclonal antibody at 1:250 dilution. DNA concentration was determined by Nanodrop (Thermo Scientific). Libraries were constructed using the KAPA Hyper Prep kit and Illumina TruSeq adapters. DNA library quality was assessed by Bioanalyzer using a High Sensitivity DNA chip (Agilent). Samples were pooled and run in a single lane on an Illumina HiSeq 4000 (Duke Sequencing and Genomics Technologies core). 50 bp single reads were assessed for quality using FastQC (www.bioinformatics.babraham.ac.uk) and adapter sequences trimmed using Trim Galore (www.bioinformatics.babraham.ac.uk) and Cutadapt (Martin, 2011). Resultant sequences were aligned to *H. salinarum* NRC-1 genome (RefSeq: NC_002607.1, NC_002608.1, NC_001869.1) using Bowtie2 (Langmead and Salzberg, 2012). Subsequent analyses were conducted in the R Bioconductor coding environment, and all associated code is freely available at https://github.com/amyschmid/cdr. Peaks were called using MOSAiCS (Chung et al., 2016) from sorted bam files with arguments: fragment length 200, bin size 200, read capping 0, analysis type IO, background estimate rMOM, signal model 2S, FDR 0.01. Peaks reproducible across two of three biological replicate samples were integrated using the DiffBind (Stark, 2011) and ChIPQC (Carroll et al., 2014) packages. Peak locations were associated with annotated genes using the IRanges Bioconductor package (Lawrence et al., 2013). Data were visualized for the figures using the R package trackViewer (Ou and Zhu, 2019). R package version numbers are given in the github repository at https://github.com/amyschmid/cdr. Raw and analyzed data are available through GEO accession GSE148065.

## Data availability

All gene expression and ChIP-seq data from this study are available to the publich through GEO accession GSE148065. Whole genome resequencing data are available via Sequence Read Archive Project PRJNA614648. Code and datasets are available on the GitHub repository https://github.com/amyschmid/cdr. All supplementary figures, tables, and movies are available on FigShare via https://doi.org/10.6084/m9.figshare.12195081.v1.

## Acknowledgments

Funding for this project was provided for NSF MCB-1651117 and MCB-1417750 to AKS, and Wellcome 203276/Z/16/Z and NIH DP2AI117923 to ECG. Special thanks to Ana Sofía Uzsoy at the North Carolina School of Science and Math for assistance with cloning and growth curve CFU quantification. We acknowledge the excellent support of Nicolas Devos at the Duke Sequencing and Genomic Technologies Core Facility and Holly Dressman at the Duke Microbiome Center Shared Resource. Thanks to Schmid lab members Rylee Hackley and Sungmin Hwang for their comments on the manuscript.

## Guide to supplementary material

Supplement can be accessed via the FigShare repository associated with this study at https://doi.org/10.6084/m9.figshare.12195081.v1.

<u>**Supplementary text.**</u> Supplementary figure legends and Supplementary Tables 4-7.

## Supplementary Figures

**Supplementary Figure S1**. The *cdrS-ftsZ2* locus is conserved across Haloarchaea, and *cdrL* is unique to halophiles.

**Supplementary Figure S2.** Cell lengths at mid-log and stationary phase.

**Supplementary Figure S3**. Micrographs of cells treated with aphidicolin.

**Supplementary Figure S4.** Design of mother machine microfluidics device.

**Supplementary Figure S5.** Growth curves for individual cells in the mother machine.

**Supplementary Figure S6.** Expression data for all genes tested by NanoString.

**Supplementary Figure S7.** CdrL-FLAG ChIP-seq data: all replicates, growth phases, and peak detection.

## Supplementary movies

**Supplementary Movie SM1.** Movie of Δ*cdrS* growing in the mother machine, then extruding out of the channel by the end of the movie.

**Supplementary Movie SM2.** Movie of Δ*ura3* with *ftsZ1*-msfGFP integrated into the chromosome.

**Supplementary Movie SM3.** Movie of Δ*ftsZ2* with *ftsZ1*-msfGFP integrated into the chromosome.

**Supplementary Movie SM4a.** Movie of Δ*cdrS* with *ftsZ1*-msfGFP integrated into the chromosome.

**Supplementary Movie SM4b.** Movie of Δ*cdrS* with *ftsZ1*-msfGFP integrated into the chromosome.

## Supplementary Tables

**Supplementary Table 1.** Archaeal genomes with UniProt identifiers for CdrS-FtsZ2 loci and FtsZ homologs with no CdrS associated.

**Supplementary Table 2.** Whole genome resequencing data for Δ*cdrS* and Δ*ftsZ2*.

**Supplementary Table 3.** Raw growth data and colony forming units (CFU) corresponding to main text Figure 2.

**Supplementary Table 4.** Nanostring raw and normalized data.

**Supplementary Table 5.** Strains used in this study.

**Supplementary Table 6.** Primers used in this study.

**Supplementary Table 7.** Plasmids used in this study

## Notes

https://doi.org/10.6084/m9.figshare.12195081.v2

https://github.com/amyschmid/cdr

### Competing Interest Statement

The authors have declared no competing interest.

